# mCSEA: Detecting subtle differentially methylated regions

**DOI:** 10.1101/293381

**Authors:** Jordi Martorell-Marugán, Víctor González-Rumayor, Pedro Carmona-Sáez

## Abstract

**Motivation:** The identification of differentially methylated regions (DMRs) among phenotypes is one of the main goals of epigenetic analysis. Although there are several methods developed to detect DMRs, most of them are focused on detecting relatively large differences in methylation levels and fail to detect moderate, but consistent, methylation changes that might be associated to complex disorders.

**Results:** We present *mCSEA*, an R package that implements a Gene Set Enrichment Analysis method to identify differentially methylated regions from Illumina 450K and EPIC array data. It is especially useful for detecting subtle, but consistent, methylation differences in complex phenotypes. mCSEA also implements functions to integrate gene expression data and to detect genes with significant correlations among methylation and gene expression patterns. Using simulated datasets, we show that *mCSEA* outperforms other tools in detecting DMRs. In addition, we applied *mCSEA* to a previously published dataset of sibling pairs discordant for intrauterine hyperglycemia exposure. We found several differentially methylated promoters in genes related to metabolic disorders like obesity and diabetes, demonstrating the potential of *mCSEA* to identify differentially methylated regions not detected by other methods.

**Availability:** *mCSEA* is freely available from the Bioconductor repository.

## Introduction

DNA methylation is by far the most studied epigenetic mark. It affects gene expression and has an important role in several disorders. Epigenome-wide association studies (EWAS) are performed to find associations between DNA methylation alterations and a given phenotype (Flanagan, 2015).

There are several methodologies to determine DNA methylation status, including high-throughput techniques such as whole-genome bisulfite sequencing (WGBS) or methylation arrays. Although WGBS is the one with highest coverage, Illumina’s BeadChip arrays (Infinium HumanMethylation450 (450K) and Infinium MethylationEPIC (EPIC)) are still much more affordable and simpler to analyze, and they are currently the most used platforms in human EWAS (Teh et al., 2016).

EWAS are usually applied to find associations between individual CpG sites and outcomes. However, methylation patterns are not usually found in isolated differentially methylated positions (DMPs). Instead of that, clusters of proximal CpGs are hypermethylated or hypomethylated (Peters et al., 2015). That is the reason why several methods have been designed to detect differentially methylated regions (DMRs) instead of individual DMPs. Some methods use predefined regions as candidates for DMRs identification (e.g. gene promoters or CpG Islands), while others do not rely on previous annotations and search *de novo* DMRs.

There are two different paradigms related to DNA methylation (Leenen et al., 2016). The first one is that, in some disorders such as cancer, regulatory regions are clearly hypermethylated or hypomethylated, with methylation differences greater than 60 % (De Smet et al., 1999). However, there is a second paradigm in which complex disorders are associated to very subtle differences in CpGs methylation, with methylation differences of 1-10 % between phenotypes (Guerrero-Bosagna et al., 2014).

Most available DMR methods have focused on detecting large methylation differences between phenotypes. In this context, they work properly and they have allowed the discovery of many epigenetic causes of several diseases (Lappalainen and Greally, 2017). However, these tools may fail to detect significant DMRs in complex diseases or heterogeneous phenotypes, where there might be small differences among methylation signals but consistent across the analyzed regions and samples. Therefore, no individual CpGs or regions may meet the threshold for statistical significance in many studies, although there may be biologically meaningful differences (see for example Bohlin et al., 2015; Chiavaroli et al., 2015; van Dongen et al., 2015; Gervin et al., 2012; Kim et al., 2017).

In addition, some of these tools average all sites in a given region, but if a significant pattern is associated to a subset of sites it may be underestimated if all sites are analyzed as a block.

This scenario motivated us to develop a new approach based on Gene-Set Enrichment analysis (GSEA) (Subramanian et al., 2005), a popular methodology for functional analysis that was specifically designed to avoid some drawbacks related such as to small statistical differences in gene expression patterns. GSEA uses a given statistical metric to rank all genes of a genome and applies a weighted Kolmogorov–Smirnov (KS) statistic (Hollander and Wolfe, 1999) to calculate an Enrichment Score (ES). Basically, ES for each set is calculated running through the entire ranked list increasing the score when a gene in the set is encountered and decreasing the score when the gene encountered is not in the analyzed set. ES of this set is the maximum difference from 0. Significance of each ES is calculated permuting the sets and recomputing ES, getting a null distribution for the ES. GSEA is capable of detecting subtle enrichments of genes in gene sets.

We adapted this approach defining gene sets as sets of CpG sites in predefined regions. We called our approach *mCSEA* (methylated CpGs Set Enrichment Analysis) and it is capable to detect subtle but consistent methylation differences in predefined genomic regions from 450K and EPIC microarrays data. The methodology has been implemented as an R package freely available in Bioconductor repository.

## Methods

### *mCSEA*’s workflow

*mCSEA* R package consists of five main functions (Figure 1). The first step of the mCSEA analysis is to rank all the CpG probes by differential methylation. This step can be done independently and the analysis start from the sorted list or using the rankProbes() function, which apply *limma* (Ritchie et al., 2015) to fit a linear model and returning the t-statistic assigned to each CpG site.

**Figure 1.**
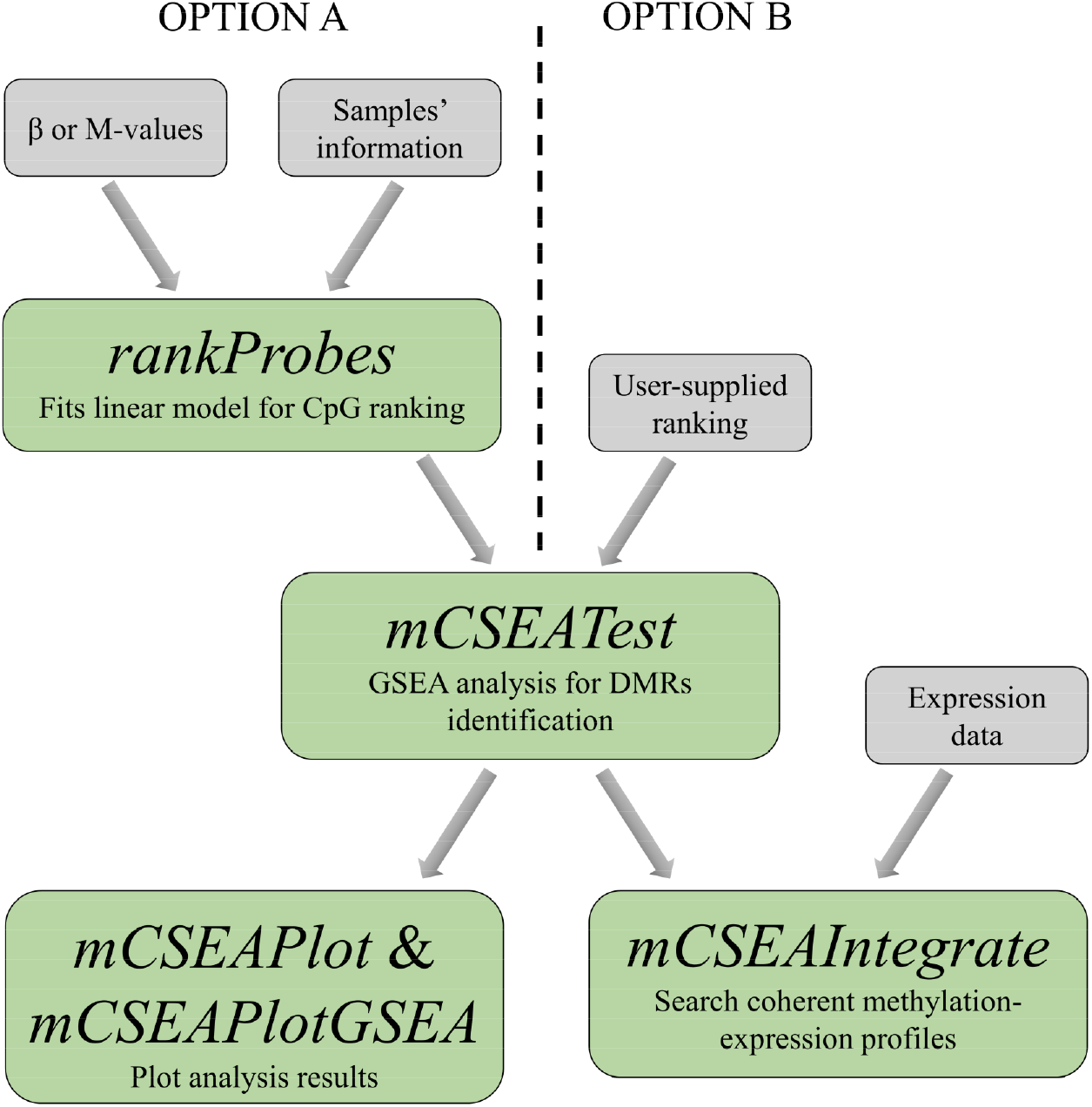
*mCSEA*’s workflow. Grey boxes are input data and green boxes are mCSEA’s functions The scheme also shows the order in which functions should be executed.

The main *mCSEA* function, mCSEATest(), evaluates the enrichment of CpG sites belonging to the same region in the top positions of the ranked list. Applying the standard GSEA method implemented in the *fgsea* package (Sergushichev, 2016) regions whose CpG sites are over-represented in the top or bottom of the list can be detected as differentially methylated. As predefined regions *mCSEA* allows users to perform analysis based on promoters, gene bodies and CGIs. In addition, the function also allows researchers to use user-defined regions in the analysis. Among other results, the users obtain a P-value for each region to be differentially methylated.

*mCSEA* package include two functions to visualize the results: mCSEAPlot() and mCSEAPlotGSEA(). The former represents methylation values of a given region in its genomic context (see Figure 3 for an example). The latter generates GSEA’s enrichment plot, showing the positions of the CpG in a determined region along the entire ranked list.

**Figure 2.**
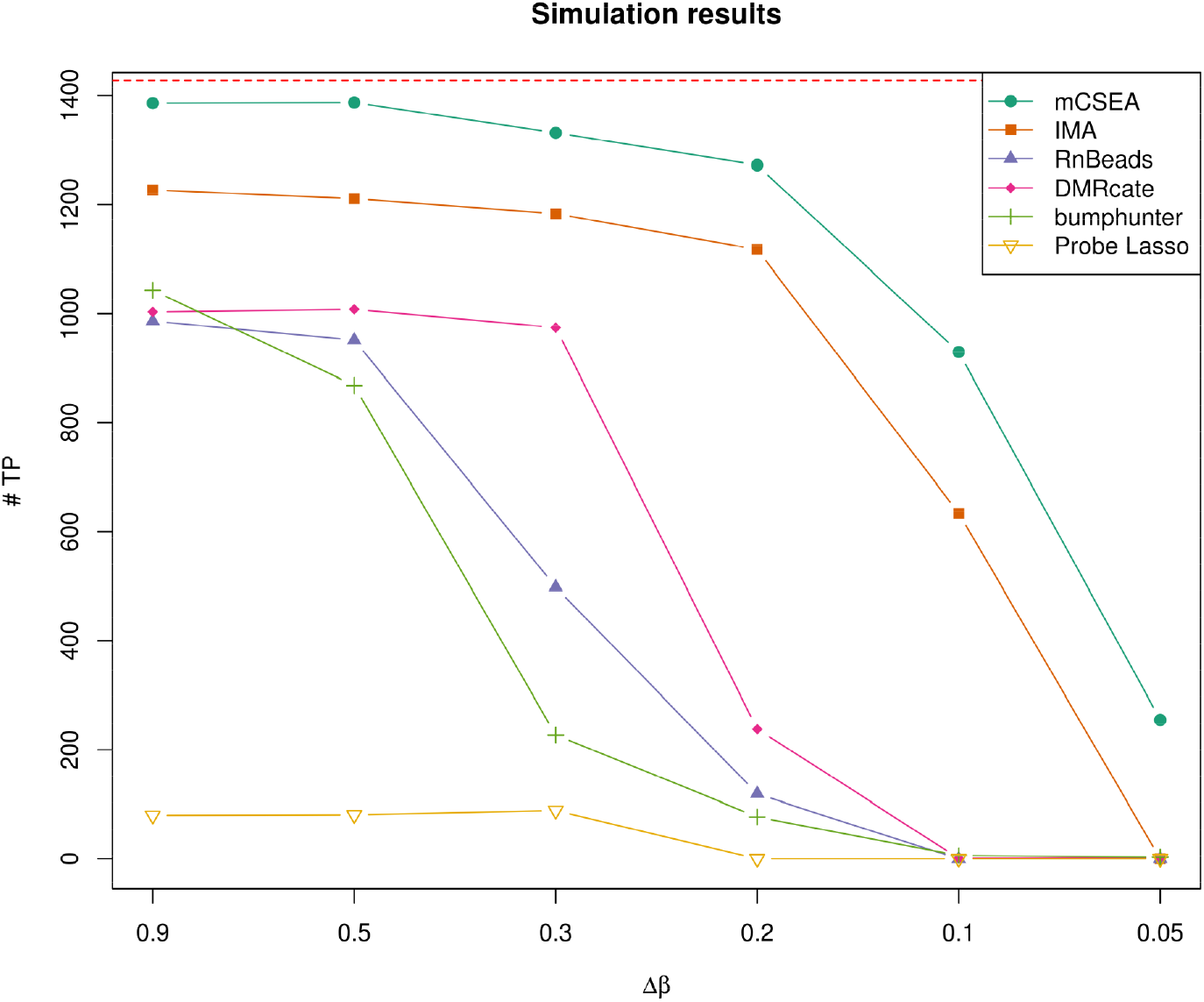
Performance in simulated data. Each line represents results from different methods. The Y-axis represents the number of TP for each Δβ. Red line represents the total number of TP included in the dataset (1428).

**Figure 3.**
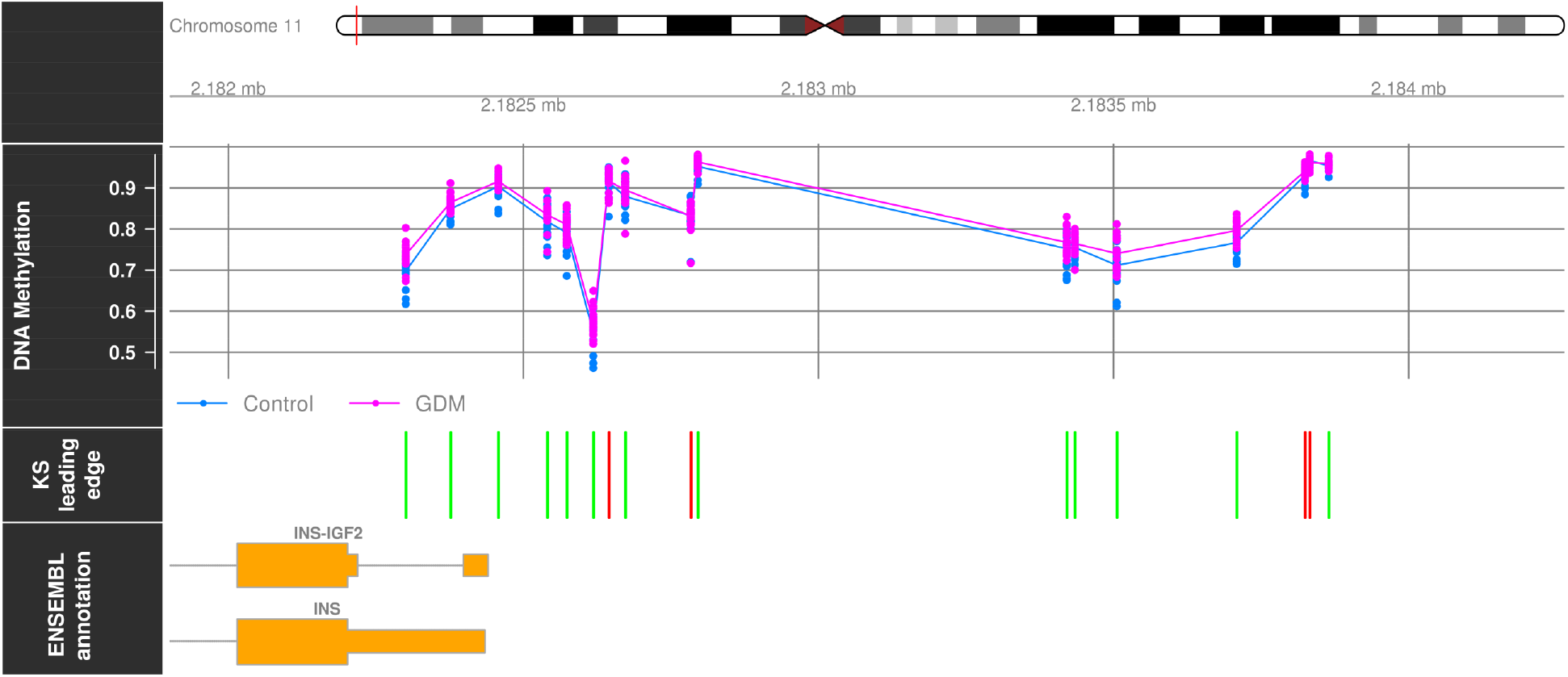
INS gene promoter methylation in GDM and control samples. Methylation is quantified with β-values. Each point represents the methylation of each sample. Lines link the mean methylation of each group. KS leading edge panel marks with green bars those CpGs contributing to the ES and with red bars the rest of them. This plot was obtained with mCSEAPlot() function, implemented in mCSEA package.

Finally, the package implements a function, mCSEAIntegrate(), which integrates gene expression data in the analysis. For that purpose, the leading edge CpGs of each region is first defined. Leading edge is the set of CpGs that contributes to the ES of the region, so these CpGs are the most differentially methylated ones. These sites are averaged for each region in each sample. Then, Pearson’s correlation coefficient is calculated between each region’s methylation and the proximal gene(s)’ expression (i.e. genes within 1500 base pairs upstream and downstream from the region). If the integration is performed with promoters, significant negative correlations are returned, due to it has been observed an inverse correlation between promoters’ methylation and gene expression (Jones and Baylin, 2002). On the contrary, if the integration is performed in gene bodies, significant positive correlations are returned instead, due to a positive correlation between gene body methylation and expression has been observed (Aran et al., 2011). If the integration is performed in CGIs, both positive and negative significant correlations are returned, due to CGIs can be located in both promoters or gene bodies.

### Methods comparison

In order to test our method, we used both simulated and real data. We simulated 450K β-values for 20 samples using the same approach as Peters et al. (Peters et al., 2015). We randomly selected 714 promoters to be hypermethylated and another 714 promoters to be hypomethylated in 10 samples (cases) compared to the other 10 (controls). Only promoters with at least 5 associated CpGs were selected. In order to not simulate an ideal situation where the entire regions are differentially methylated, we randomly chose the first or the second half of the regions to be hyper- or hypomethylated, what is more similar to the expected in a real dataset. We simulated datasets with a β-value mode differences among phenotypes (Δβ) ranging from 0.9 to 0.05 across promoter CpG sites. We compared *mCSEA*’s performance with state-of-the-art solutions, both predefined (*IMA* (Wang et al., 2012) and *RnBeads* (Assenov et al., 2014)) and *de novo* (*DMRcate* (Peters et al., 2015), *bumphunter* (Jaffe et al., 2012), and *Probe Lasso* (Butcher and Beck, 2015)) algorithms. *IMA* package uses as input raw idat files and not β-values matrix. Therefore, to compare its approach using the simulated data we implemented the method that is applied by *IMA*, that is to calculate the median of the methylation values for each predefined region and to apply *limma* to these averaged values. We did not include *COHCAP* package (Warden et al., 2013) due to it restricts the analysis to CGIs. For all methods we used default parameters with the exceptions compiled in Supplementary Table 1.

**Table 1.**
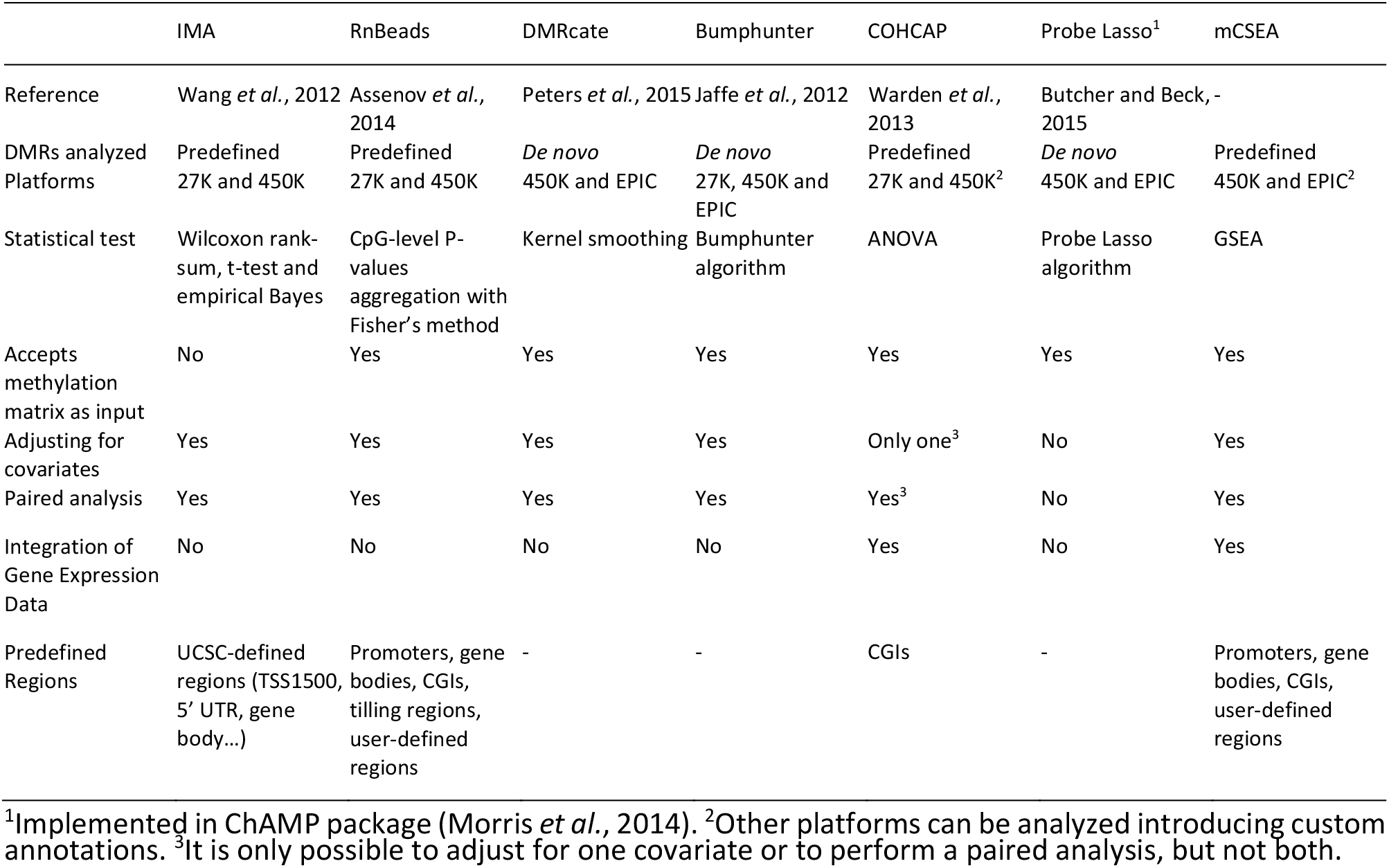
Comparison of available R packages for DMRs analysis using Illumina’s microarray data.

All results were considered significant using P-value adjusted by false discovery rate (FDR) < 0.05 threshold. For *IMA* and *RnBeads*, we searched for DMRs in promoter regions and we considered as true positives (TP) those promoters annotated with the actual differentially methylated promoters, and as false positives (FP) the called regions not annotated with the actual DMRs. For the rest of the methods, due to they return *de novo* DMRs, we considered as TP those actual DMRs overlapping at least one called region, and as FP the called regions not overlapping any actual DMR. For all methods we considered as false negatives (FN) the actual DMRs not called by the corresponding method.

For each method and Δβ we calculated the sensitivity (Equation 1) and the precision or positive predictive value (PPV) (Equation 2).

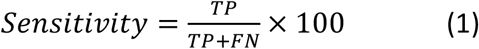

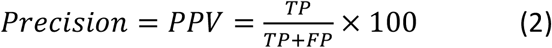

We also tested the performance of the proposed method in the methylation datasets previously published by Kim et al., 2017. This dataset contains Illumina 450K methylation data from 18 sibling pairs discordant for intrauterine exposure to maternal gestational diabetes mellitus (GDM). This data is publicly available from GEO database (GEO ID: GSE102177). We reanalyzed the data with *IMA*, *RnBeads*, *DMRcate*, *Probe Lasso*, *bumphunter* and *mCSEA*. We selected these methods because all of them are popular tools for DMRs analysis and allow complex experimental designs with paired samples and covariates, as was our case. Although Probe Lasso does not allow this kind of designs, we modified its code to include this feature.

## Results

### Comparison of DMRs analysis packages

We performed a functional comparison of *mCSEA* and the most popular R packages used to DMRs analysis from Illumina microarrays data (Table 1). An essential function of this kind of software is the capability to analyze data from complex experimental designs, due to methylation data is very sensitive to environmental factors (Marsit, 2015) and it is important to take into account sex, age, ethnicity and other cofounding factors. In addition, some experiments require a paired analysis (e.g. when normal and cancer cells are extracted from the same patient). *mCSEA* can handle with both, covariates adjusting and paired analysis. Other important features analyzed were the type of regions that can be included in the analysis and the capacity of integrating gene expression data.

### Simulated data results

We calculated the number TP, FP and FN returned by each tested method for each Δβ situation, in addition to sensitivity and PPV (Supplementary Table 2). As can be noted in Figure 2, *mCSEA* yielded the best performance detecting methylation differences ranging from Δβ=0.9 and Δβ=0.2 and it mainly outperformed the rest of methods when the methylation differences were especially small (0.1 and 0.05). In addition, *mCSEA* returns a low number of FP, resulting in a high PPV for all Δβ. Only *DMRcate* overcomes mCSEA in PPV, but at the cost of having a significantly lower sensitivity for all Δβ.

### DMRs in maternal diabetes exposure discordant siblings

To demonstrate the mCSEA’s functionality, we analyzed the data reported by (Kim et al., 2017). This is a methylation dataset from child sibling pairs: one of the siblings was exposed to maternal diabetes during their gestation, while the other was not. This intrauterine hyperglycemia exposure is associated to an increased risk of obesity and diabetes. Authors collected data from discordant siblings for maternal diabetes exposure in order to get insight into possible epigenetic aberrations in the exposed sibling. Methylation differences in such type of experiment were expected to be very subtle and, in fact, original authors did not find any significant result (FDR < 0.05).

In our reanalysis, *DMRcate* and *Probe Lasso* did not return any significant DMR. These methods applied *limma* to detect significant DMPs and call DMRs based on them. Although they work properly when methylation differences are high they did not reveal any significant result for slight methylation differences.

*RnBeads* is also based on *limma* for detecting DMRs, but it combines the results by region types (promoters, CGI, and so on) aggregating the P-values obtained by the linear modelling, so even if there are not any significant DMPs, *RnBeads* is potentially capable to find significant DMRs. However, this was not the case. This method did not return any significant DMR (FDR < 0.05).

*IMA* approach did not return any significant DMR neither.

*Bumphunter* yielded one significant DMR (FDR = 0.03, FWER = 0.01) located at the promoter of SDHAP3 pseudogene. Up to our knowledge, there is not any known relationship between SDHAP3 and development or metabolic disorders.

*mCSEA* yielded 1055 significant DMRs (FDR < 0.05) in gene promoters: 228 hypermethylated and 827 hypomethylated promoters in cases compared to controls (Supplementary Table 3).

To assess the biological significance of these results, we performed an enrichment analysis using Enrichr (Chen et al., 2013). The most significant enriched pathway in KEGG database (Kanehisa and Goto, 2000) is “Maturity onset diabetes of the young” (hsa04950) pathway (adjusted P-value = 0.0011) (Supplementary Table 4). This pathway is related with a type of diabetes characterized to appear in patients younger than 25 years old and to be non-insulin dependent. Promoter regions of nine out of the twenty-six genes associated to this pathway were identified as significantly differential methylated regions, including PDX1, FOXA2, PAX6 or INS. The INS gene, which we found to be hypermethylated in cases, is an important one that has been previously associated to diabetes in several works and it has been reported as a silenced gene with a fully methylated promoter associated to diabetes development (Yang et al., 2011). In addition, it has been observed that high levels of glucose increase the INS methylation level (Yang et al., 2011), so this hypermethylation could be induced during gestation. Methylation differences in INS promoter between children exposed and non-exposed to intrauterine hyperglycemia are subtle, but consistent across all CpG sites of the promoter (Figure 3). Such small methylation difference is the cause why this DMR remains undetected by all the other tested methods. The same may be occurring in many other genomic regions.

On the other hand, the most significant enriched pathway from OMIM Disease database is obesity (adjusted P-value = 0.0085) (Supplementary Table 5). Eight out of fifteen genes related to this disease contained significant DMRs, including UCP1, UCP3, GHRL or PCSK1. So, we found methylation alterations in genes related to diabetes and obesity, the two main diseases associated to intrauterine GDM exposure.

## Conclusions

Here we present *mCSEA*, a novel R package for predefined DMRs detection based on GSEA method. We compared *mCSEA* with the most widely used methods to detect DMRs. Our method outperformed the rest of solutions for detecting small methylation differences in the simulated dataset. It is especially remarkable the capability of *mCSEA* to find DMRs even with the methylation difference as small as 0.05 between groups, but consistent along a relatively large region. We reanalyzed a previously published dataset, obtaining barely no significant results with other methods. However, *mCSEA* yielded several significant DMRs in promoters for genes associated to relevant biological pathways.

We think that *mCSEA* will provide researchers with a useful tool to detect DMRs in datasets from complex diseases in which the methylation differences among phenotypes are small but consistent.

## Funding

JMM is partially funded by Ministerio de Economía, Industria y Competitividad. This work is partially funded by Consejería de Salud, Junta de Andalucía (Grant PI-0152-2017).

